# Dissecting Metabolic Landscape of Alveolar Macrophage

**DOI:** 10.1101/2023.09.08.556783

**Authors:** Sunayana Malla, Karuna Anna Sajeevan, Bibek Acharya, Ratul Chowdhury, Rajib Saha

## Abstract

The highly plastic nature of Alveolar Macrophage (AM) plays a crucial role in the defense against inhaled particulates and pathogens in the lungs. Depending upon the signal, AM acquires either classically activated M1 phenotype or alternatively activated M2 phenotype. These phenotypes have specific functions and unique metabolic traits such as upregulated glycolysis and pentose phosphate pathway in M1 phase and enhanced oxidative phosphorylation and tricarboxylic acid cycle during M2 phase that help maintain the sterility of the lungs. In this study, we investigate the metabolic shift in the activated phases of AM (M1 and M2 phase) and highlight the roles of pathways other than the typical players of central carbon metabolism. Pathogenesis is a complex and elongated process where the heightened requirement for energy is matched by metabolic shifts that supplement immune response and maintain homeostasis. The first step of pathogenesis is fever; however, analyzing the role of physical parameters such as temperature is challenging. Here, we observe the effect of an increase in temperature on pathways such as glycolysis, pentose phosphate pathway, oxidative phosphorylation, tricarboxylic acid cycle, amino acid metabolism, and leukotriene metabolism. We report the role of temperature as a catalyst to the immune response of the cell. The activity of pathways such as pyruvate metabolism, arachidonic acid metabolism, chondroitin/heparan sulfate biosynthesis, and heparan sulfate degradation are found to be important driving forces in the M1/M2 phenotype. We have also identified a list of 34 reactions such as nitric oxide production from arginine and the conversion of glycogenin to UDP which play major roles in the metabolic models and prompt the shift of the M2 phenotype to M1 and vice versa. In future, these reactions could further be probed as major contributors in designing effective therapeutic targets against severe respiratory diseases.

**Author Summary:** Alveolar macrophage (AM) is highly plastic in nature and has a wide range of functions including invasion/killing of bacteria to maintaining the homeostasis in the lungs. The regulatory mechanism involved in the alveolar macrophage polarization is essential to fight against severe respiratory conditions (pathogens and particulates). Over the years, experiments on mouse/rat models have been used to draw insightful inferences. However, recent advances have highlighted the lack of transmission from non-human models to successful *in vivo* human experiments. Hence using genome-scale metabolic (GSM) models to understand the unique metabolic traits of human alveolar macrophages and comprehend the complex metabolic underpinnings that govern the polarization can lead to novel therapeutic strategies. The GSM models of AMs thus far, has not incorporated the activated phases of AM. Here, we aim to exhaustively dissect the metabolic landscape and capabilities of AM in its healthy and activated stages. We carefully explore the changes in reaction fluxes under each of the conditions to understand the role and function of all the pathways with special attention to pathways away from central carbon metabolism. Understanding the characteristics of each phase of AM has applications that could help improve the therapeutic approaches against respiratory conditions.

## Introduction

Alveolar Macrophages (AM) are the first line of defense against respiratory pathogens and are highly plastic in nature(1). Depending upon the interactions with pathogens, AMs can be polarized into several subsets(1). The two main subsets known are: classically activated or pro-inflammatory (M1) macrophages and alternatively activated or anti-inflammatory (M2) macrophages(2,3). M1 macrophages respond to microbial factors like Lipopolysaccharide (LPS) and Th1 pro-inflammatory cytokines that play a significant role in bacterial killing and recruitment of other immune cells(4–7). M2 macrophages, on the other hand, can be induced by interleukin 4(IL-4) that promotes anti-inflammatory activities such as resolution to inflammation and repairing of the damaged cells(8). Both the phenotypes are marked by their unique metabolic niches - such as enhanced Glycolysis and Pentose Phosphate Pathway (PPP) in the M1 phase and Oxidative Phosphorylation (OXPHOS), Tricarboxylic Acid Cycle (TCA), and Fatty Acid Oxidation (FAO) in M2 phase(9). The reprogramming of AMs towards M1 or M2 phenotype is contingent upon specific signaling molecules including IFN-γ (type II interferon), LPS, IL-4, and immune complexes (Ic)(10). The resultant phenotype is influenced by the prevailing physiological demands, such as pathogenic bacterial killing or tissue restoration, that determine the sequential progression of the phenotype development(8,11,12). In infected lung tissue, AMs are first polarized to the M1 phenotype and later to M2 phenotype for a healthy immune response(13). However, alterations in the interaction could be catastrophic to the cell(1). The shift in phenotypes of AM plays an important role in regulating the body’s immune response and metabolism(14,15). Yet, the regulatory mechanisms governing the polarization are not completely understood and little attention has been dedicated to the role of physical parameters such as temperature (1–3).

The thermal component of fever and its effect on inflammation is one of the most poorly understood aspects of pathogenesis (16,17). Fever is the first immune response against respiratory pathogens such as tuberculosis, influenza, and SARS-CoV2. The increase in core temperature enables the formation of pro-inflammatory M1 cells to combat the invading bacteria(18). However, it has also been reported that fever enables the M2 phenotypic behavior to maintain homeostasis in the lungs(16). Hence, understanding the relationship between fever and macrophage polarization is crucial to understand the regulation of macrophage function ^15^. Experimental analysis on a mouse model by increasing core temperature to 39.5 ^O^C, was found to have significant positive effect on the modulation of macrophage function (19). However, several documented examples provide evidence on the disparity between the experimental conclusions in rodent studies and humans(20). Due to the challenges associated with human experimental studies, systems biology approaches can be utilized for reconstructing context-specific (i.e., M1 or M2 phase or elevated temperature) Genome-Scale Metabolic (GSM) models. Systems biology has been proven to be very useful for pragmatic modelling and theoretical exploration of complex biological systems(21). By the integration of high-throughput omics data (metabolomics, proteomics, or transcriptomics), the reconstruction of human GSM models of pancreatic cancer(22), tuberculosis(23), obesity and diabetes’s(24), neurodegenerative diseases(25) has led to discovery of novel therapeutic targets and better understanding of the metabolic shifts.

GSM models are computational context specific (species, cells, tissue, etc.) knowledgebases capable of dissecting systemic metabolic phenomena(26). A GSM model contains all annotated metabolic reactions and pathways within a biological system^26^. Using Flux Balance Analysis (FBA) and Flux Variability Analysis (FVA) the fluxes of the reactions can be predicted for a given condition/timepoint. The study by Gelbach et al, on M1 and M2 subtypes of human colorectal cancer cells via generation of GSM models marks a significant step towards understanding and manipulating the polarization mechanism of macrophages(27). However, it is also important to explore the metabolic network and capabilities of M1 and M2 phenotypes in tissue-specific macrophages (e.g., AM) where they show unique behaviors and patterns. The early GSM of AM was curated by Bordbar et al, from the global human model, Recon1(28), in 2010 and was further used to model tuberculosis infected AM(23). However, the model did not consider other potential states (e.g., M1 and M2 phase of AM) which is why it is highly critical to dissect the metabolic interactions in AM in its activated state with pathogens to fully understand the progression of the disease. However, thus far not much attention has been devoted to the contextualization of the AM model to represent M1 and M2 state and the effects of physical parameters such as temperature(4,17,29).

In this study, three context specific GSM models were generated by integrating transcriptomics data of healthy AM, and its activated phases: M1 and M2. **Figure 1** shows the overview of steps involved in generation, curation, and analysis of the three GSM models. Integration techniques such as iMAT (30) and E-flux(31) were used for GSM model reconstruction from Human1(32). The metabolic models were further curated by using a tool called OptExpand (inhouse tool, currently unpublished) that can be used to identify and resolve Thermodynamically Infeasible Cycles (TICs). These models were used to investigate the altered metabolism of activated AM (M1 and M2 phase) compared to that of healthy AM and compare M1 and M2 phases with each other as well. The healthy AM model was validated by reproducing experimentally reported rate of ATP production and Nitric Oxide production(33,34). The comparison of the flux ranges showed enhancement in Glycolysis, Pentose Phosphate Pathway (PPP), and shift in Tricarboxylic Acid Cycle (TCA) to accumulate succinate and itaconate in M1 phase. M2 phase reaction fluxes show upregulation of Oxidative Phosphorylation (OXPHOS), uninterrupted TCA cycle, and upregulated Fatty Acid Synthesis (FAS). These observed metabolic shifts in M1 and M2 phases are in accordance with the previously reported evidence from literature(34). In addition, the metabolic pathway was found to be more active as the temperature increases from 38°C to 41°C. Some unique characteristics of pathways such as Pyruvate Metabolism, Glycolysis, Carnitine Shuttle (mitochondria) pathway, and Bile Acid Synthesis (BAS) were further explored to understand specific nature of each activated state, highlighting the role of Chondroitin/Heparan Biosynthesis and Heparan Sulphate degradation as a potential point of manipulation in M1/M2 balance. Going forward, the context specific activated AM GSM models will be used to study interaction with respiratory pathogens. In addition, models of a system of immune cells such as AM, Neutrophils, and Mast cells, could be developed to analyze the intercellular interactions during pathogenesis.

**Figure 1:**
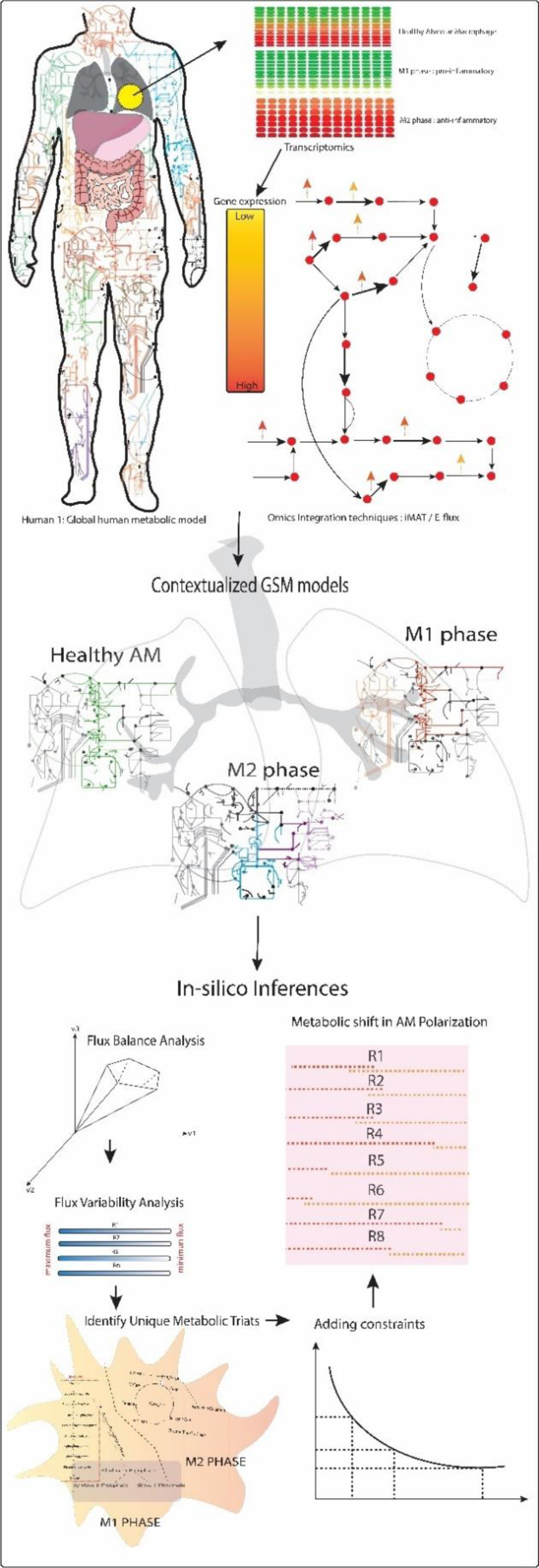
Schematic of the workflow for the generation of healthy and activated Alveolar macrophages and the steps involved in analysis of the metabolic shift during polarization.

## Results and Discussion

### Metabolic Model Reconstruction of Alveolar Macrophage (AM) Metabolism

Genome scale metabolic (GSM) models provide an improved understanding on the metabolic basis of different biological processes and have been widely used for biomedical applications(26). These sophisticated cellular systems of metabolic reactions in conjunction with their corresponding genes and enzymes provide novel insight into the initiation and progression of diseases(35). The interconnection between the genes, metabolites, and reactions are converted into mathematical representation and fluxes are predicted by performing computational flux analysis such as flux balance analysis (FBA) and flux variability analysis (FVA)(36,37). The latest global human metabolic reconstruction, Human1(32) is an extensively curated representation of human metabolism that combines two parallel lineages, namely Recon(28,38,39) and Human metabolic Reaction (HMR)(40). With 20% higher total reactions, 33% higher metabolites, and higher mass balance than those of any available human reconstructions, Humn1 has successfully been used to reconstruct cell-specific GSMs for liver, liver cancer, blood, blood cancer etc.(32). Hence, this standardized model allows the convenient integration of omics data to reconstruct AM specific metabolic model.

In this work, metabolic reconstruction of healthy AM, M1 phase, and M2 phase was obtained by integrating gene expression values of metabolic genes(9,41,42) onto the Human1 model. These transcriptomic profiles were acquired from GEO databases (GSE8823, GSE40885, and GSE41649 for AM, M1 phase and M2 phase, respectively). Among various methods available to integrate omics data, iMAT(30), a switch approach, indicates the presence/absence of a specific reaction depending on the relevant gene(s) having higher expression levels at a specific condition. On the other hand, a valve approach such as E-flux(31), uses gene expression levels to control the flux of the corresponding reactions. The healthy AM model obtained upon the implementation of iMAT consists of 4,554 reactions (governed by 2,173 metabolic genes) and 3,967 metabolites (2,003 unique) distributed across eight intracellular compartments (Extracellular, Peroxisome, Mitochondria, Cytosol, Lysosome, Endoplasmic reticulum, Golgi apparatus, Inner mitochondria, and Nucleus); while the model generated by E-flux consists of 8,073 reactions and 5,380 metabolites (2,823 unique) across these eight compartments with the same number of metabolic genes. Since both these approaches employ different fundamental assumptions (as mentioned above) and are usually more successful in different applications(43). The sensitivity of iMAT approach to user defined threshold typically leads to higher number of reactions to be omitted or sometimes leads to exclusion of important reactions from the pruned model. In our case, this resulted in a version of the pruned model capable of producing biomass, but it failed to include important pathways such as NO production, glycerolipid metabolism, heme synthesis, and porphyrin metabolism. The E-flux-generated model, on the other hand, consists of comparatively higher number of reactions, metabolites, and included all the important pathways mentioned earlier. **Figure 2** shows the distribution of active reactions in the important AM pathways. Additional information on each model can be found in supplementary files. Similar observations were obtained while implementing iMAT and E-flux with the expression values of 2,951 and 2,390 metabolic genes to reconstruct GSM models for M1 and M2 phases respectively (additional information on the gene expression values, distribution of pathways, reactions, and metabolites for all the models are available in supplementary files). The GSM model developed upon the E-flux method for M1 phase consists of 7,986 reactions, and 5,602 (2,821 unique) metabolites and the similar model of M2 phase model consists of 7,884 reactions and 5,936 (2,969 unique) metabolites. On the other hand, the pruned models obtained from iMAT implementation were significantly smaller with important biological pathways missing. Hence, for the purpose of our study, E-flux was able to incorporate all the important AM pathways, active reactions, and metabolites (with a higher number of unique metabolites). This allowed us to exhaustively investigate the metabolic shift occurring during polarization.

**Figure 2:**
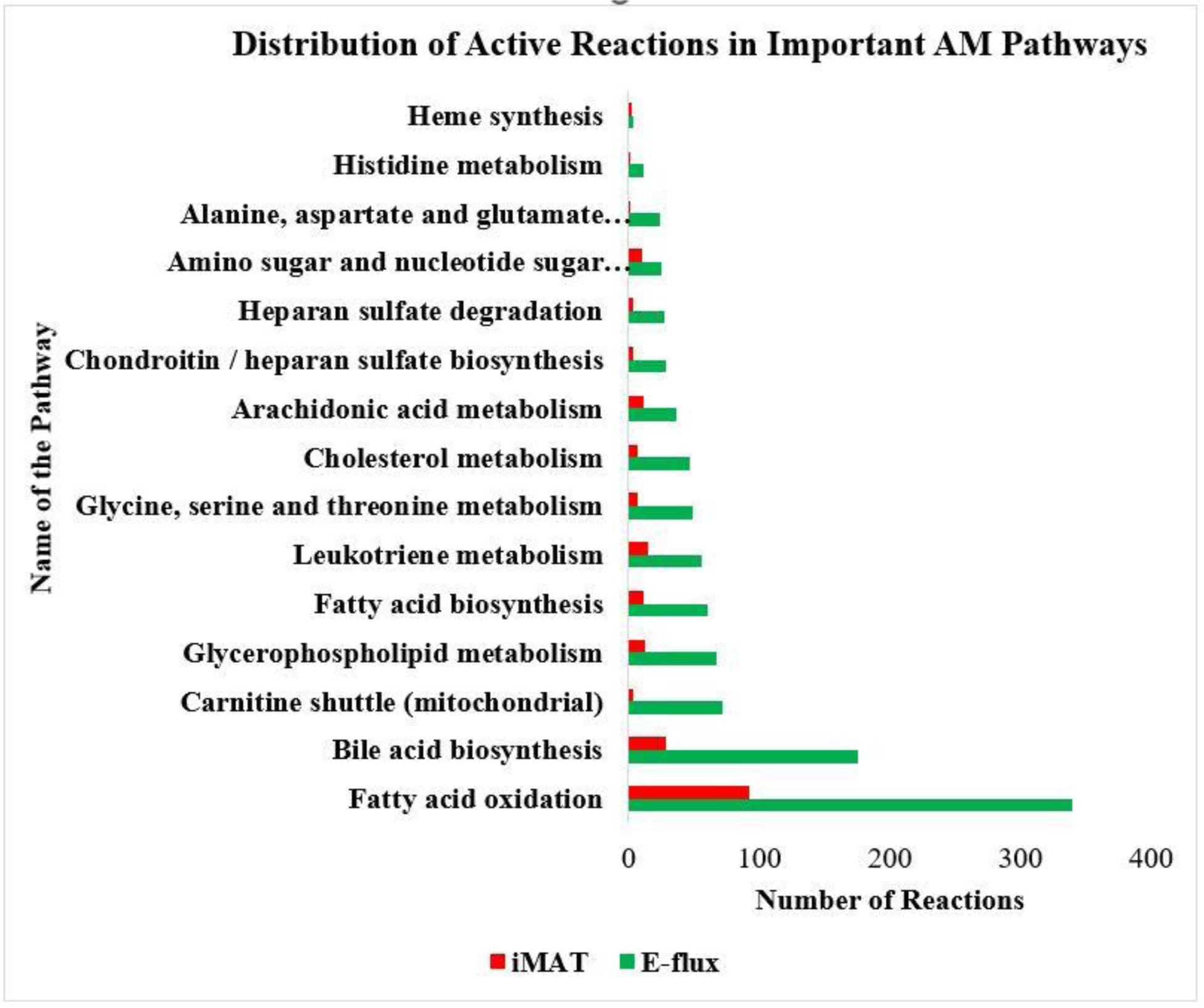
The distribution of active reactions in the context-specific generated models via iMAT and E-flux. The figure highlights the inclusive properties of the applied approaches.

The models were next validated to ensure their ability to simulate biologically significant processes by reproducing important metabolite production rates and characteristic behavior as reported in literature. FBA is used to optimize the production rates of important healthy AM metabolites while maintaining the maximal level of biomass (i.e., 0.03 h^-1^). AM are tissue resident macrophages that populate the lung environment during birth and last for the lifespan of the individuals typically(23). Since AMs do not readily multiply, the biomass function comprises of mainly cellular maintenance requirements such as proteins, lipids, DNA repair, ATP maintenance and RNA turnover(23). The model was optimized for ATP production and NO production that yielded the flux of 0.6 mm/h/g cell DW and 0.03 mm/h/g cell DW respectively. These *in-silico* values were very close to the values of 0.71 mmol/h/g cell DW and 0.037 mmol/h/g cell DW, respectively, as obtained from *in vitro* experiment(33,34). Hence, the healthy AM model obtained via E-flux algorithm is capable of reproducing important experimentally reported production rates.

Similar to healthy AM model, the activated phase models should also be able to recapitulate the relevant metabolic reprogramming. To this end, with the help of the GSM of healthy AM as the base, the M1/M2 phase reaction fluxes were compared that generated four possible scenarios: overlap of flux with increase, complete overlap with decrease, partial overlap with increase, and no overlap with increase in forward direction. As reported in literature for M1 phase, we observed increased activity (complete overlap and partial overlap) for glycolysis pathway(9). Pentose Phosphate Pathway (PPP) shows 48% of the reactions have increased fluxes which include important reactions such as formation of Ribulose 5-phosphate and its conversion to Ribose 5-phosphate with production of NADPH. This is a crucial step in energy production during M1 phase(9). We also found the increased production of succinate and itaconate in the TCA cycle which limits the formation of precursors that aid oxidative phosphorylation (OXPHOS) and electron transport chain (ETC)(9,44). Similarly, the comparison of the reaction fluxes between M2 phase and healthy AM showed increased activities in OXPHOS, fatty acid oxidation (FAO), and TCA cycle without extra accumulation of succinate and itaconate metabolites. FAO impairs the anti-inflammatory responses and helps OXPHOS increase the production of ATP through TCA cycle. These metabolic traits are well supported by literature(9,45) and thus establish the credibility of our context-specific GSM models of the activated phases.

### A Response to Fever: Increase in Temperature

Fever is the highly evolved systematic inflammatory response that is not limited to the site of infection but affects the whole body(16). The heat of fever is reported to supplement the performance of immune cells by increasing stress on pathogens and infected cells(17). However, the advantages of fever in different conditions are still not clear. For instance, lowering of temperature could be more beneficial than its increase in cases of extreme inflammation(18). The study of effects of physical parameters such as temperature in human physiology is very important but is extremely challenging to investigate due to numerous limitations such as higher cost to conduct *in vivo* or *in vitro* studies to gather human cell data(46,47). Here, we study the fluctuation in the thermodynamic feasibility of reactions and pathways in response to change in temperature by calculating change in Gibbs free energy (ΔG) of reactions and Max/Min Driving Force (MDF) of the pathways(48). Starting from the core temperature of human body (37°C), the analysis is completed till 40°C, beyond which fever is considered fatal(49).

Equilibrator was used to calculate the standard Gibbs free energy of formation (Δ*f*G°) for the reactions of interest(50). Equilibrator so far is equipped to calculate the standard Gibbs free energy of reactions with KEGG(51) and BIGG IDs(52), hence limiting the number of full pathways we could analyze. In addition, due to the lack of available AM and temperature specific metabolomics, the metabolite concentration ranges were set to be 1 nM to 10 mM which is the typical metabolite range for a biological system capable of capturing adequate cellular physiology(48). The response of pathways such as glycolysis, OXPHOS, PPP, TCA cycle, amino sugar and nucleotide sugar metabolism, leukotriene metabolism and some amino acid biosynthesis pathways (proline/alanine, and arginine biosynthesis) were investigated by calculating MDF. The change in Gibbs free energy at different temperatures gave us insight into the thermodynamic feasibility of each reaction and pathway during the progression of fever (increase in temperature). A steady increase in MDF for all the pathways was observed indicating that the increase in temperature as positive catalyst for metabolism. The study detailed and specific response of each reaction/pathway to temperature will be possible with further advancement in human metabolomics experimental data generation. The calculated MDF for all of the above pathways was above 10KJ/mol, indicating the thermodynamic favorability of the pathways as the temperature increases. With the low driving force (less than 3 KJ/mol) the reactions are found to be heavily dependent on kinetic parameters such as enzyme concentration and turnover rate (i.e., kcat)(53). However, the dependence of the reaction rate on the kinetics decreases with the increase in driving force. It is reported that with a driving force of 10KJ/mol or higher. the reactions occur in forward direction with negligible flux in reverse direction(48). In **Figure 3a**, the reaction with the maximum ΔG in glycolysis pathway is highlighted with orange. Similarly, the reactions from TCA cycle (**Figure 3b**) with maximum Gibbs free energy are also highlighted. Both the reactions in the pathways are key steps for ATP production which are found to be more feasible with the increase in temperature. The change occurring in leukotriene metabolism in comparison to Δ*f*G° is also shown in **Figure 3c**, which indicates the increasing thermodynamic feasibility of reactions in Leukotriene metabolism with the increase in temperature. Full details on all the other pathways mentioned above are present in the supplementary files. Assuming the set metabolite concentration is favorable and the Δ*f*G° values are accurate, as the temperature increases the thermodynamic feasibility of the pathways also increases, the reactions occur in the forward direction spontaneously with less enzymatic effort. This phenomenon ultimately supports the biological need for enhanced metabolic activities to illicit immune response in the cell. We observed small but noticeable changes in MDF values and ΔG values for all the reactions and pathways. With further advances on availability of experimental data, in future we can further explore the extent of effect of temperature.

**Figure 3:**
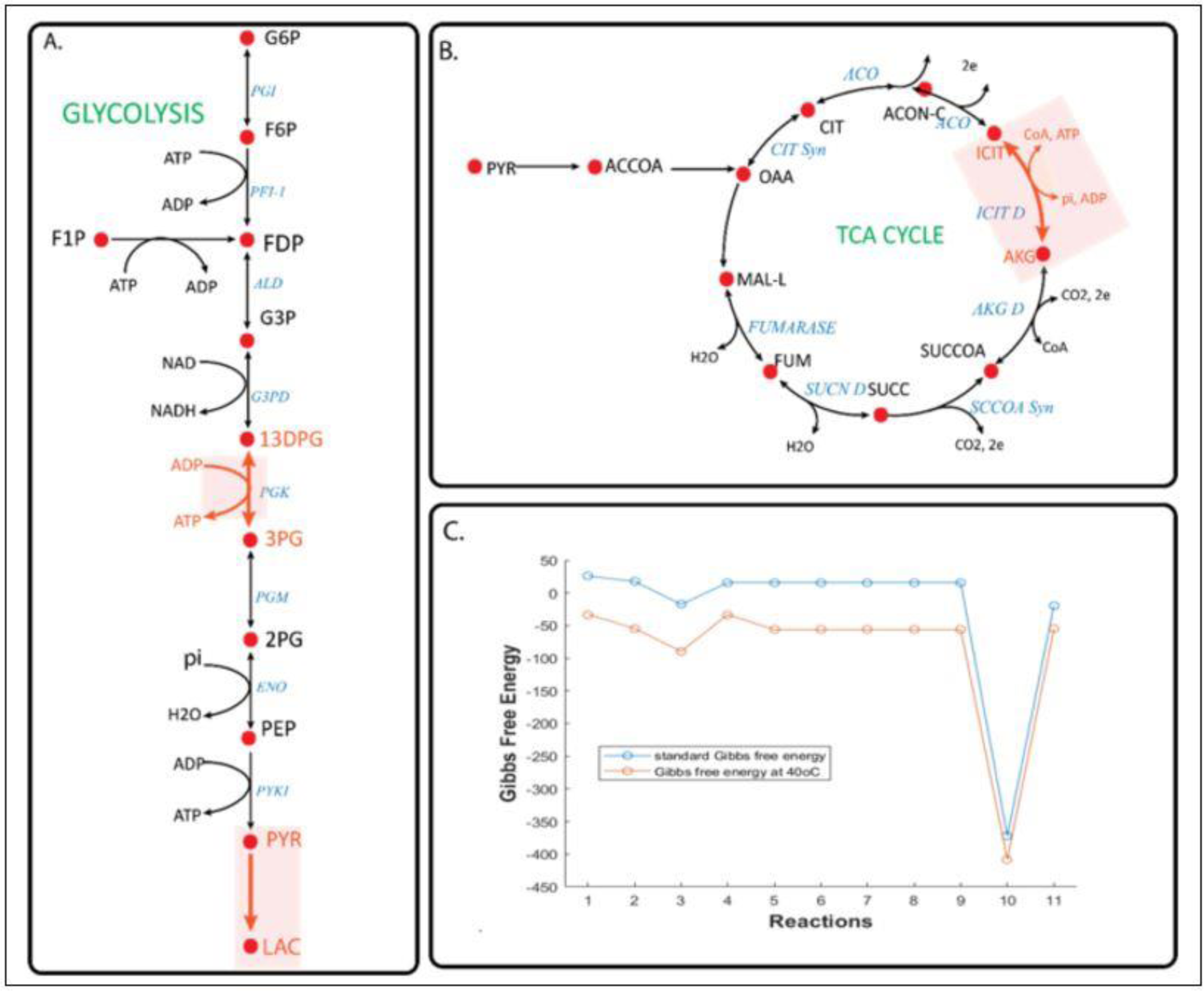
Part A and B showcase the maximum Gibbs free energy of a reaction in the major pathway such as Glycolysis and TCA cycle. The final figure C shows the change for the reactions in Leukotriene metabolism.

The enzymes govern the direction and rate of reactions at a molecular level. The change in Gibbs free energy for a reaction is directly associated with the enzyme turnover rate also known as kcat. To study the change in the enzyme turnover rate during the macrophage polarization, four enzymes catalyzing reactions from the pathways mentioned above were selected. To determine suitable values of kcat, deep learning pipeline DLkcat(54) trained on SABIO-RK(55) database were first used. Non-trivial differences in these predicted values prompted us to put together a new enzyme structure-aware method of calculation for kcat. Due to paucity of exhaustive experimental kcat measurements, we shortlisted four enzymes that were part of our prior analysis and had literature evidence towards inflammatory or anti-inflammatory responses(56–60). The proteins, GRPHR (glyoxylate and hydroxypyruvate reductase). OCD1 (ornithine decarboxylase 1), GLS (glutaminase), and GNE (glucosamine (UDP-N-acetyl)-2-epimerase/N- acetylmannosamine kinase) catalyze the reactions, glyoxalate to glycolate in mitochondria, conversion of ornithine to putrescine and carbon dioxide in extracellular matrix, conversion of glutamine to glutamate and ammonia, and N-acetyl-D-mannosamine to N-acetylmannosamine-6- phosphate in cytoplasm respectively. Next, enzyme turnover rate (kcat) and saturation (K) were used to explain the change in concentration of these enzymes (E) at different values of maximum velocity (Vmax). Vmax represents the maximum possible flux for the reactions in each of the activated phases obtained from FVA. All the calculations for determining E can be found in supplemental files. We found that enzyme concentration and the enzyme saturation relation differ for each gene with the change in Vmax, indicating the difference in their role during inflammation/anti-inflammation responses. The enzyme concentration was higher at all saturation points in M1 phase for GRHPR and ODC1 gene, while the GLS concentration was high for M2 phase and GNE enzyme concentration was found similar for both the phases. The activity of each of these enzymes provides insight into the metabolic reprogramming occurring in AM while acquiring the desired phenotype. For example, the presence of glyoxalate at various concentrations has been associated with inflammation and diseases which are governed by GRHPR(56). Similarly, the conversion of ornithine to putrescine which occurs in presence of OCD1 is a key *in vivo* biomarker for higher parasite survivals(57,58). In order to further explore and understand the metabolic shift at pathways and reactions levels, we next investigated the individual reactions fluxes of activated AM GSM models with Healthy AM.

### *De novo* Metabolic Reprogramming in AM Polarization Mechanism

AM in the lungs act as the first line of defense against respiratory pathogens since these phagocytize pollutants and pathogens that act as a trigger to activate an innate immune response(14). M1 and M2 macrophages acquire distinct phenotypes which are usually driven by different stimuli. M1 macrophages are stimulated by LPS and IFN-γ which enable the production of pro-inflammatory cytokines such as IL-1, IL-12, IL-23, ROS(59). On the other hand, M2 phase is stimulated by IL-4 or IL-13 which promotes anti-inflammatory cytokines releasing IL- 10(60–63). Expanding our attention beyond the typical players of central carbon metabolism, we see the activity of pathways such as pyruvate metabolism, arachidonic acid metabolism, chondroitin/heparan sulphate biosynthesis, and heparan sulphate (HS) degradation to be major contributors to inflammatory or anti-inflammatory responses. Despite the increasing interest in AM polarization and their unique contribution to the progression and suppression of diseases, not much attention has been given to these pathways (pyruvate metabolism, arachidonic acid metabolism, chondroitin/heparan sulphate biosynthesis, and Heparan Sulphate (HS) degradation) in lung pathogenesis. **Figure 4** shows the flux distribution in seven different pathways that play important role during AM polarization.

**Figure 4:**
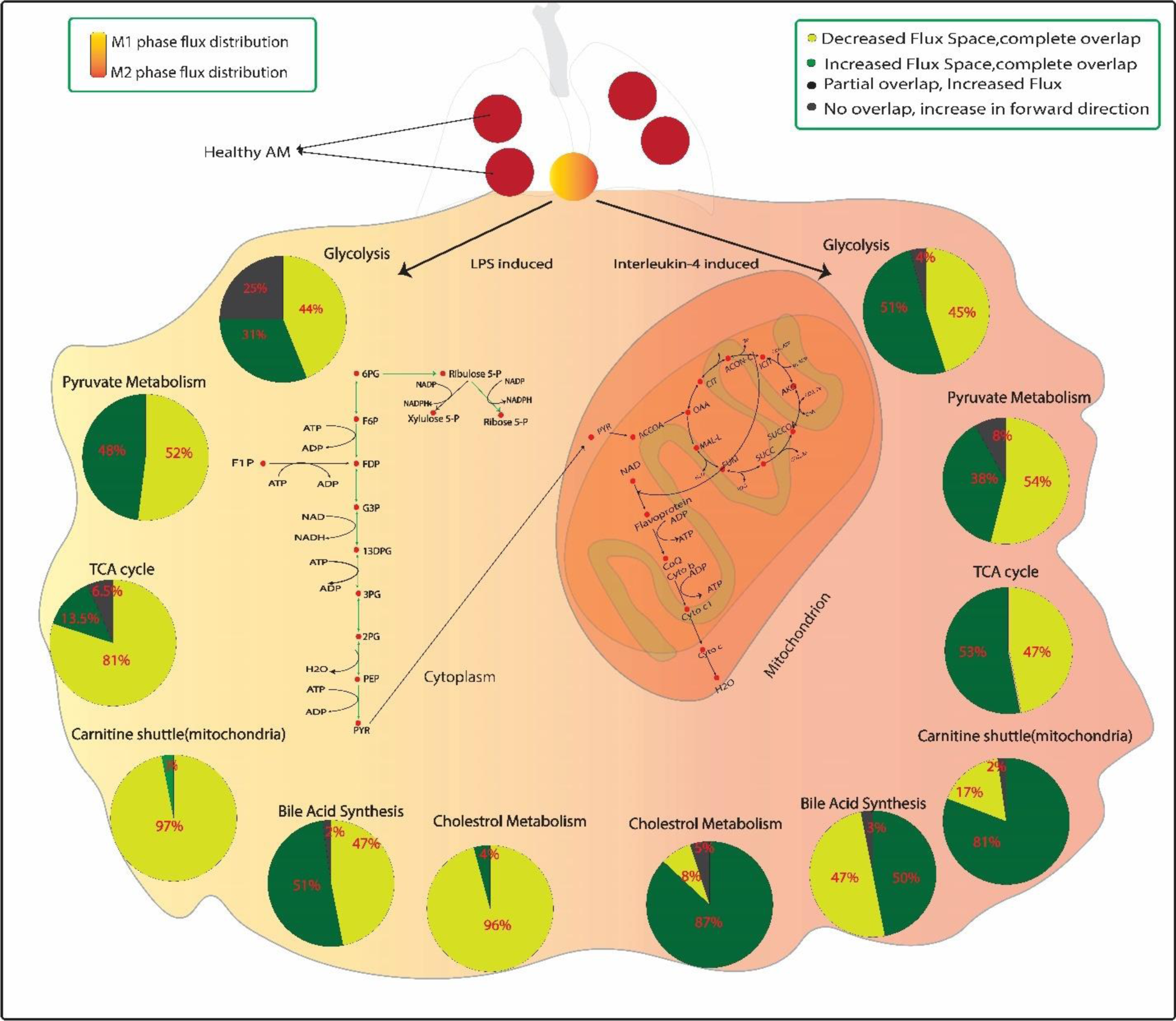
Alveolar Macrophage acquires unique metabolic characteristics depending upon the phenotype. In the M1 phase, the reactions of glycolysis are enhanced which are highlighted by the green arrows and the PPP reactions which is a major contributor for NAPH production is also enhanced. Similarly, the pathways highlighted by yellow arrows in M2 phase are found to be enhanced. Each pie chart represents metabolic reprogramming of AM in the specific pathway in either M1 phase or M2 phase. Each component of the pie chart represents one of the four categories as color coded in the figure. The associated percentage in the pie chart represents the percentage of overall reactions of a specific pathway falling into each of the categories.

As pathogens invade the lung microenvironment, the polarization shifts firstly toward M1 phenotype development(61). M2 phase is usually described as the anti-inflammatory stage where the cell mainly focuses on remodeling and tissue repair(62). For a long time, it was believed that M1 and M2 phases were drastically different both phenotypically and functionally(62). However, recent interest in the ambiguous nature of M2 phase has led to the discovery that M2 phase cells can be further divided into M2a, M2b, M2c, and M2d subtypes and each of these subtypes has its unique functions ranging from tissue repair to phagocytosis and some level of pathogen defense as well(64,65). To understand the metabolic shift in AM when it acquires M1/M2 phenotypes, the flux range of a specific reaction in the activated phase was compared with the flux range in healthy AM.

To fight the invading microorganism, cells increase toxicity to reduce the chances of survival. The M1 model exhibited higher activity in bile acid synthesis and arachidonic acid metabolism. These pathways increase the toxicity in the cell hence limiting the growth of pathogens(66–68). The reactions, 5,6-Ep-15S-HETE and 5,15-DiHETE from arachidonic acid metabolism had increased flux space in comparison to healthy state. These reactions are involved in the formation of oxygenated polyunsaturated fatty acids called oxylipins. Oxylipins play a very important role in the regulation of inflammation and the formation of other important leukotriene metabolites such as LTA4(69–71). Additionally, the activity of glycolysis was not found to be completely inhibited in M2 phase despite OXPHOS and TCA cycle showing distinctly enhanced fluxes. The nature of glycolysis activity has been a point of debate in AM polarization and with recent findings(65) that indicate M2d subtype delineates proinflammatory responses, we propose all M2 subtype population may not exhibit inhibited glycolysis activity based on our *in-silico* predictions. However, most of the energy does come from OXPHOS in M2 subtypes. Our M2 model, as stated before, was able to capture the enhanced fluxes in OXPHOS and TCA cycle and predicts over 90% of reactions to be enhanced in carnitine shuttle (mitochondria) pathway. With the help of carnitine shuttle (mitochondria) pathway, long-chain fatty acids that are impermeable to mitochondrial membranes are migrated into the matrix for β-oxidation and energy production(72). Hence, upregulated activity of carnitine shuttle pathway is unique to M2 phase as we did not observe similar activity in M1 phase. In fact, the flux analysis of M1 phase with healthy state showed us over 80% of the reactions to have inhibited fluxes. **Figure 5** portrays the complex regulatory pathways in AM polarization.

**Figure 5:**
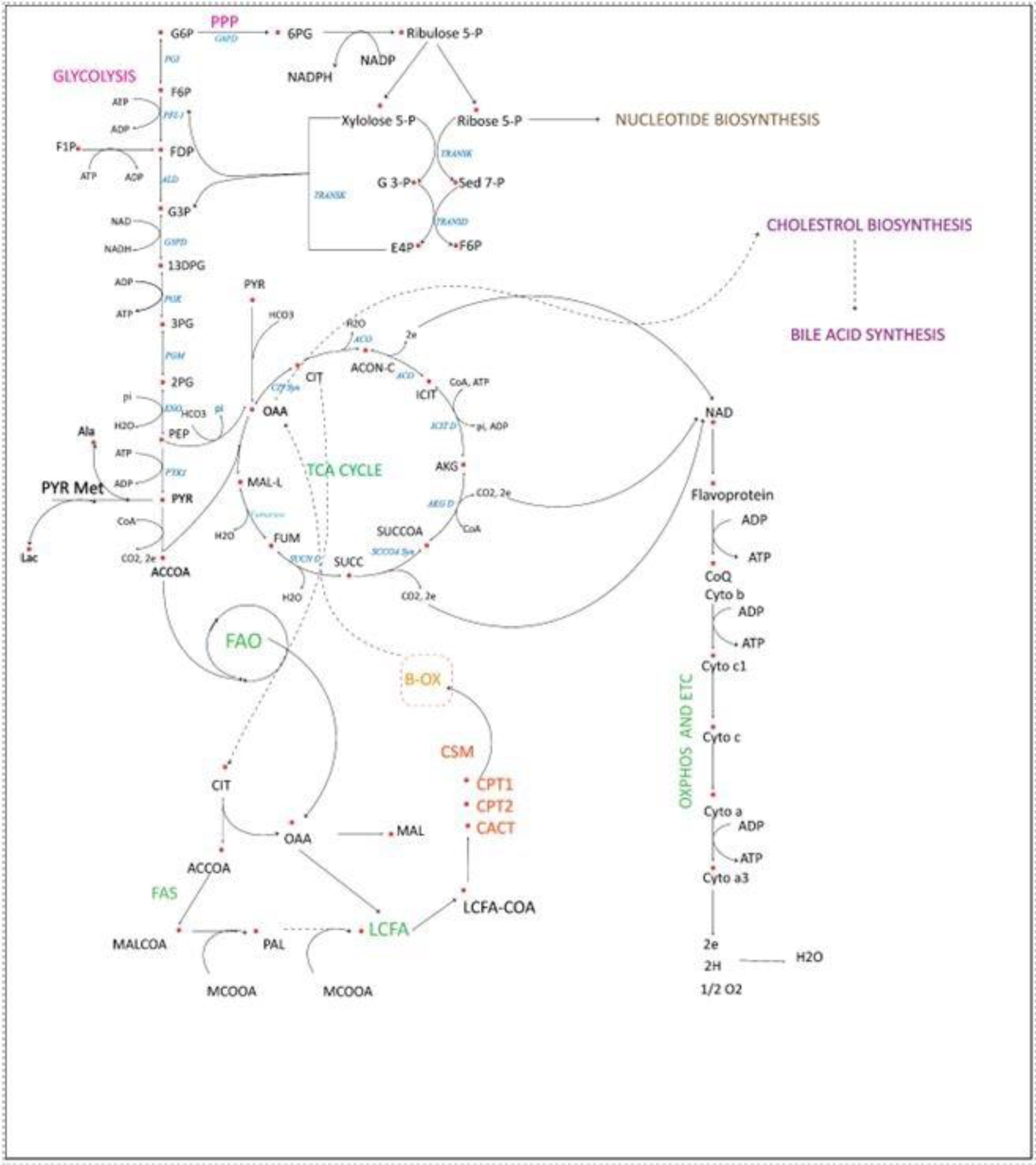
Important AM pathways. Glycolysis, TCA cycle, and OXPHOS play major roles in energy production with the help of pathways such as the carnitine shuttle (mitochondria), which shows enhanced activity during the anti-inflammatory phase. On the contrary, Bile Acid Synthesis, and Arachidonic Acid Metabolism are heightened to induce acidic conditions to minimize pathogen survival. Pyruvate Metabolism play key roles in the immune response of the cell.

Upon exploring further, the activity of Chondroitin/heparan sulfate biosynthesis and HS degradation were found to be specific to each phase as well. These pathways either contribute to the formation or degradation of an important metabolite called Heparan sulphate. The formation of heparan sulfate is a crucial step for the recruitment, adhesion, crawling, and transmigration of leukocytes from the circulation to the site of inflammation(73). And the mechanism related to the initiation of the inflammatory response is Chondroitin/heparan sulfate biosynthesis(74,75). The role of formation and degradation of HS has been a topic of interest during lung injury and inflammation; however, it has not been highly studied(75). We found all the active reactions were enhanced in M1 phase from Chondroitin/heparan sulphate biosynthesis while the Heparan Sulphate (HS) degradation was found to be enhanced during M2 phase when compared to healthy AM. Hence, the *in-silico* activity of Chondroitin/heparan biosynthesis suggests enhanced inflammatory response while HS degradation is related to the versatile function in M2 phase or slightly inhibited inflammatory response. We further explored the activity of these pathways between M1 and M2 phase and expanded on it in the next section.

Furthermore, the activity of pathways such as pyruvate metabolism was expected to be enhanced as pyruvate is a key mediator in cellular metabolism. In addition to the typical glycolysis to TCA cycle pathway, pyruvate can also be derived from lactate and from amino acids such as arginine (76). Despite being such a key modulator, the pyruvate metabolism, as a whole, was found to be inhibited in M1 as well as M2 phase from our models with respect to the flux activity in healthy AM. We found that the reactions contributing to the direct formation of pyruvate were mostly inhibited in both the activated macrophages. A study by Abusalamah(77) suggests that incorporating pyruvate as sodium pyruvate in growth media for macrophages inhibited immune response of the cell and also had positive impact on the bacterial growth(77). We found the overall activity of pyruvate metabolism was inhibited in the sense that excess pyruvate production is inhibited. The key reaction such as PEP to pyruvate at the end of glycolysis and pyruvate to OAA at the beginning of TCA maintained high fluxes but other reactions that contribute to pyruvate through different mechanisms were inhibited. This indicates that not only pyruvate metabolite but the whole metabolism plays a crucial role during polarization.

### Delicate Balance between M1 and M2 Cells

Pathogenesis in the lungs is usually marked by an influx of M1 cells which later turn into M2 cells(78). However, interaction with certain pathogens inhibits or promotes the development of certain phenotypes to ensure the survival of the virus. For example, the interaction between tuberculosis and AM is sometimes reported to promote M2 cells in comparison to M1, and reports on progression of cancer cells also mention the positive role of the M2 phenotype(79–81). The imbalance in the M1/M2 cells can be deleterious to the lungs that can cause prolonged and unwanted inflammation in the absence of the process that shut it down. In addition, without the necessary inflammation, AMs cannot effectively activate other immune cells to fight invading organisms(82). Hence it is very important for AM to shift towards the phenotype which is best suited to fight the invading pathogens. The unique metabolic shift in M1 and M2 phenotypes with respect to healthy AM is crucial to understand the diseased state, however, it is equally important to understand the rebuttal mechanism of theAM(83). One way to study how AM maintains defense against pathogens could be by understanding the balance between M1 and M2 phenotypes and the possible reaction activities that can promote the shift from one phenotype to another. With increased interest in the role of AM as the first line of defense against respiratory pathogens, a lot of attention has been given to the signaling pathways(84–86). Manipulating signals to the cell has yielded promising results in obtaining M2 cells from M1 and vice versa, especially in rodents and *in vitro* studies(87,88). However, not much attention has been given to reactions and pathways in human cells(23). Identifying the specific pathways (reactions and metabolites) through GSM models could be a huge step forward to obtain highly effective therapeutic targets and shift the development of cells toward the desired phenotype(89). We compared the fluxes obtained from FVA with the M1 phase as the base condition. The activity of M2 phase fluxes was categorized into five different conditions (namely, complete overlap: widened flux space, complete overlap: shrunk flux space, partial overlap: increase, no overlap: definite increase in forward direction, and no overlap: definite increase in reverse direction).

Earlier, we noted that despite being a key intermediatory metabolite, the metabolic shift in AM resulted in limited pyruvate production in both M1 and M2 phases with respect to healthy AM. By comparing the M1 fluxes with M2 phase fluxes, we observed all the reactions contributing to pyruvate production in the cytoplasm were inhibited (complete overlap, shrunk flux space) whereas the mitochondrial reactions are enhanced (complete overlap, widened flux space). In **Figure 6**, it is shown that in the cytoplasm only one reaction (malate to pyruvate) has enhanced fluxes with higher fluxes toward the production of D-Lactate. On the contrary, enhanced fluxes were observed in multiple reactions that lead to pyruvate with more L-lactate production. The pyruvate produced in mitochondria is directly used up for OAA which promotes OXPHOS and TCA cycle. And lactate plays an important role in the maintenance of acid-base balance in the cell and plays a crucial role in the maintenance and resolution of inflammation(90,91). Hence, the *in-silico* flux activity suggests pyruvate metabolism is a key player to ensure proper inflammatory response and anti-inflammatory responses. Further experimental studies in human alveolar macrophage could establish not only pyruvate metabolite as an important factor but also recognize the regulation of pyruvate metabolism as a key step in pathogenesis. In addition to the metabolites from pyruvate metabolism, glycogen was also found to play an important role in regulating the inflammatory/anti-inflammatory responses of each phenotype. We observed the category with “definite increase in forward and reverse direction” consisted of mainly reactions related to glycogen.

**Figure 6:**
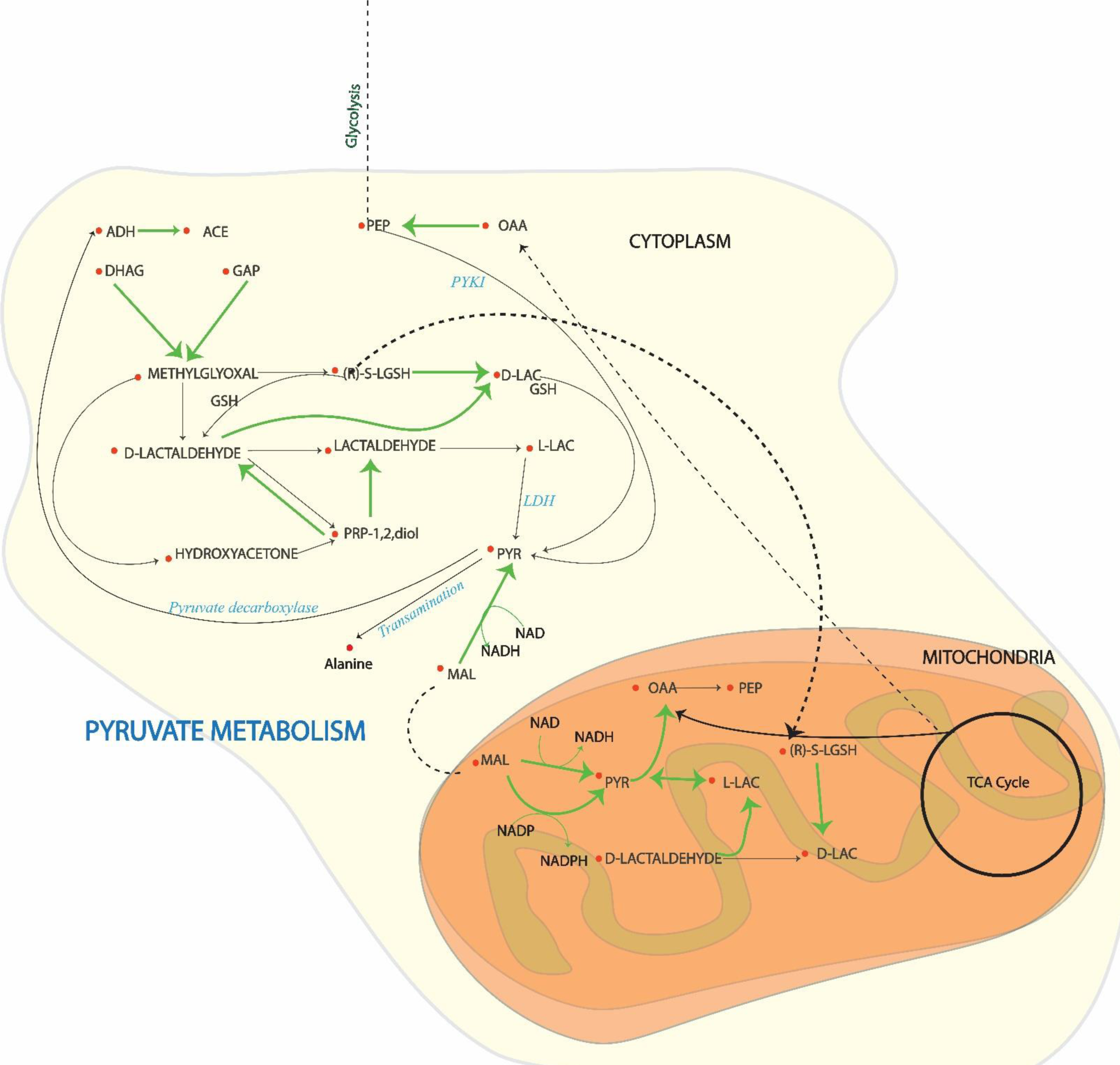
Pyruvate Metabolism activity in activate phase M2 when compared to M1 phase. The reactions indicated by green arrow are enhanced in the M2 phase.

Glycogen is one of the major sources of nutrients in AM and it has been linked to important roles in inflammation as well as maintaining surfactant production ensuring the correct lung expansion during breathing(92,93). The upregulated activity of this category of reactions starts from the production of glycogenin G8 from glycogenin in cytoplasm and finally results in the production of glycogenin G4G4. Although the importance of glycogenin as an enzyme for the regulation of glycogen is talked about but it’s true potential is not fully explored(92). The *in-silico* activity of these reactions highlights the role of glycogen as a potential target to manipulate the shift of the phenotypes. To further examine the role of these reactions, FBA and FVA were used to constrain the model by turning off the reactions completely or by limiting the flux of each reaction. We found that by turning off the reaction, UTP: alpha-D-galactose-1-phosphate uridylyltransferase, the solution becomes infeasible. This reaction is responsible for the conversion of UTP and alpha-D-galactose-1-phosphate to UDP-galactose in the cytoplasm. UDP-galactose is an essential metabolite for building galactose-containing proteins and fats that play crucial roles related to chemical signaling, building chemical structures, transporting molecules, and producing energy(94).

To be able to force M2 cells behave like M1 and vice versa, we added multiple constraints on the GSM models. Human AM consists of complex and large regulatory networks and the generated GSM models closely resemble the three states of the cell. We know certain metabolites and reaction activities are very distinct to each phenotype such as higher ATP production in the M2 phase, and the phenomena of NO production via inducible nitric oxide synthetase (iNOS) in the M1 phase or via arginase in the M2 phase. Additional constraints in the flux range of reactions that are distinctly higher in the M2 phase in comparison to the M1 phase were also incorporated. Understanding the complex nature of biological systems that consists of numerous alternative pathways, we were able to shortlist 34 reactions which when constrained in the M2 phase gives us a modified flux range that is closer to the M1 phase. The list of reactions and the flux constraint information are available in supplementary files. The shift in certain reactions is very distinct and closely resembles the M1 phase, for example, the conversion of glucose-1-phosphate to glucose-6-phosphate (initial step of glycolysis) shifted from -10 to 18 mm/h/g cell DW to -5.9 to 18.6 mm/h/g cell DW. Also, an important step of PPP in the M1 phase, that is the conversion of ribose-1-phosphate to ribose-5-phosphate, which is also a major contributor to the production of NADPH is shifted. The high-level activity of pathways such as chondroitin/heparan sulfate biosynthesis and bile acid synthesis were also observed. Constraining the exchange of ATP, production of NO from arginine, conversion of glycogenin to UDP allowed the shift to M2 from M1. The details on the flux constraints added are available on the supplemental files.

We calculated the distance between M1 and M2 GSM model and M1 and Modified M2 GSM model by applying approaches as described in Methods and Materials. The obtained Jaccard similarity index for M1 vs M2 and M1 vs modified M2 is 0.0108 and 0.0305, and the average Jaccard distance calculated was found to be 0.73 and 0.71 respectively. Jaccard distance is the measure of dissimilarity between two sets and has been used to analyze the heterogeneity of bacteria and Human cells GSM models(95). However, there have been concerns regarding the correct representation of flux modulations via Jaccard distance. The calculated values for Jaccard similarity index and the average Jaccard distances do not show a vast difference between the two conditions. Hence, we further explored Hausdorff distance, a highly versatile and robust approach in flux modulation analysis. It gives a comprehensive measure of the dissimilarity or similarity between different conditions(96). The average Hausdorff distance between M1 and M2 phase is 53.9806 while the average between the M1 phase and modified M2 was found to be 37.9455. Additionally, the sum of the distances for both cases was found to be 4.4329 × 10^5^ and 3.1161×10^5^ respectively. The Hausdorff distance is a unitless measure elucidating the overall dissimilarity between two sets of data and hence the values obtained clearly indicate the decrease in distance when modifications are introduced to the M2 GSM model. Moreover, the t-SNE (t-distributed Stochastic Neighbor Embedding) plot was used to visualize the structure of each data set representing the M1 phase, M2 phase, and Modified M2 to observe the overall shift from M2 to M1 phenotype as shown in **Figure 7**.

**Figure 7:**
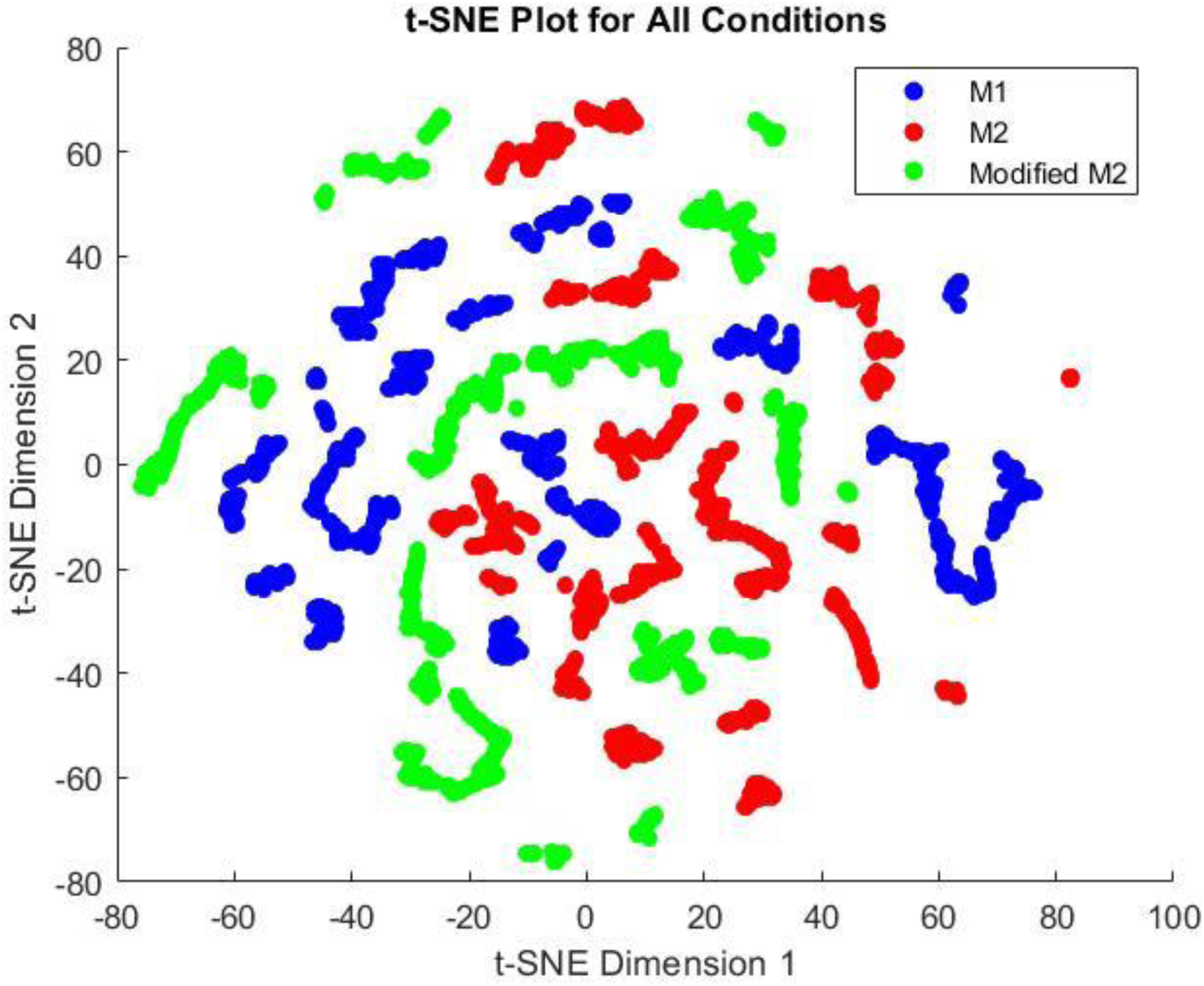
t-SNE plot visualizing the M1 phenotype, M2 phenotype, and the modified M2 phenotype represented by blue, red, and green, respectively. A distinct shift in the M2 phenotype can be observed when compared to modified M2. Modified M2 is the representation of resulting fluxes after the addition of constraints in the M2 GSM model that now resembles the M1 phenotype more closely than normal M2.

## Conclusion

The role of AM in health and disease has been a point of debate for a long time and through continuous effort from the scientific community, it has been possible to establish the versatile nature this cell exhibits(83). From the inflammatory responses to activation of other immune cells to maintaining lung homeostasis, the interest in the role of AM has opened doors to many interesting possibilities(2). Most experimental studies are limited to rodents and murine species but understanding the behavior of these cells in humans is crucial(97). Hence, Genome-Scale Metabolic models are a great initiative to understand and predict the behavior of cells under stress (pathogen invasion, nutrient deficiency, high/low temperature, etc.) conditions(21). Here, we successfully reconstructed three GSM models of healthy AM and its activated phases (M1 and M2) by integrating transcriptomics into the Human1 model. These models are capable of reproducing key biological phenomena of each state of AM cell. Here, we highlight the role of fever in the very early stages of pathogenesis as generally a positive reinforcement to the cell. Due to lack of temperature-specific transcriptomics and metabolite concentration ranges, we used a biologically feasible range (1 µM to 10 mM) of a cell and observed a steady increase in MDF of pathways. The exhaustive analysis of metabolic shifts in the activated phases with respect to healthy AM and flux comparison of M2 phase vs M1 phase was conducted. Hence, the reactions responsible for the production of oxylipins, that are directly responsible for eliciting inflammatory responses and chondroitin/heparan sulfate biosynthesis to be enhanced in the M1 phase whereas the M2 phase showed upregulated carnitine shuttle (mitochondria), and Heparan Sulphate degradation. Pyruvate Metabolism showed similar downregulated behavior in both phases when compared to healthy AM but when compared among themselves, pyruvate metabolism seemed to favor the production of OAA in the M2 phase which helps the high activity of OXPHOS, TCA cycle, and ETC in mitochondria. By understanding the key metabolic shifts, we were able to identify 34 reactions that include ATP production, NO production, glycogenin regulation, and galactose regulation reactions (such as conversion of alpha-D-galactose-1-phosphate to UDP-galactose in cytosol) which when relaxed or constrained, shift M1 phenotype to M2 and vice versa in some capacity. We can further refine and make this shift more prominent with the incorporation of metabolomics and or proteomics data. In the absence of such information, manipulating the reaction fluxes resulted in new flux ranges in M2 that have high correlations with the M1 phase and vice versa. In future, experimental validation could lead to pathways such as Heparan Sulphate degradation, Pyruvate Metabolism, and reactions involving glycogenin and galactose regulation as key players in pathogenesis. By using these GSM models, the interaction of pathogens with the AM in healthy state and activated state can be exhaustively explored. With further incorporation of human specific metabolomics/proteomics datasets when available, the temperature associated behavior of the cells could also be further studied.

## Methods and Materials

### Transcriptomics Data Processing

An exhaustive literature search was conducted to identify the appropriate set of transcriptomics data which included the transcriptomic profiles of healthy non-smokers(41), AM induced by Lipopolysaccharides (LPS) and interleukin-4 resembling M1 phase(42) and M2 phase(98), respectively. The data obtained were used as input for Gene Set Enrichment Analysis (GSEA) tool(99). GSEA is a tool that is used for pathway analysis based on the transcriptomic state of the cells and was used to compare the pathway activity of healthy AM with the M1 phase, healthy AM with the M2 phase, and M1 phase vs. M2 phase. In this process, genes are ranked based on the correlation between their expression and the class distinction using any suitable metric. GSEA calculates the enrichment score (ES) and its significance level using p-values(99). The output from the GSEA run generated lists of enriched pathways for the M1 and M2 phase. An exhaustive literature search was conducted to identify the appropriate set of transcriptomics data which included the transcriptomic profiles of healthy non-smokers’(41) AM induced by Lipopolysaccharides (LPS) and interleukin-4 resembling M1 phase(42) and M2 phase(98), respectively. The data obtained were used as input for Gene Set Enrichment Analysis (GSEA) tool(99). GSEA is a tool that is used for pathway analysis based on the transcriptomic state of the cells and was used to compare the pathway activity of healthy AM with the M1 phase, healthy AM with the M2 phase, and M1 phase vs. M2 phase. In this process, genes are ranked based on the correlation between their expression and the class distinction using any suitable metric. GSEA calculates the enrichment score (ES) and its significance level using p-values(99). The output from the GSEA run generated lists of enriched pathways for the M1 and M2 phase that mainly focused on signaling pathways.

Using the raw data set, we deduced a list of genes that were also present in the Human1 metabolic model. The list of genes for healthy AM, M1 phase, and M2 phase were 2,173, 2,951, and 2,390 respectively. The expression values of these genes were integrated into Human1 model to reconstruct models of healthy AM and its activated phases.

### GSM Model Reconstruction

The transcriptomics data obtained for each of the phenotypes of AM was integrated into Human1, a global human metabolic reconstruction consisting of 13,417 reactions, 10,138 metabolites (4,164 unique), and 3625 genes(32). Three context-specific AM metabolic reconstructions were obtained by implementing both switch and valve approaches of omics integration. Among various methods available in both the categories of switch and valve approach, iMAT and E-flux were used in our study. iMAT (integrative metabolic analysis tool) is an optimization-based program that can be used to integrate the available omics data with GSM network models for the prediction of metabolic fluxes(30). The modified version of iMAT was used where instead of classifying the overall reactions into three categories (highly expressed, lowly expressed, and moderately expressed), the reactions were divided as either highly expressed or lowly expressed with the biomass precursors always included in the highly expressed set. The formulation was constructed in such a way that all the reactions from the highly expressed set were always made active and the minimum number from the lowly expressed reaction set was added to obtain the specified objective. This resulted in a pruned mode significantly smaller than the original human model with reactions, and metabolites specific to AM and its activated stages. On the other hand, E-flux only requires the change of the upper bound and lower bound on each reaction depending on the gene expression level(31). The bounds are normalized to range between -1000 to 1000. The forward reactions consisted of a lower bound of 0 and a unique upper bound according to the gene expression levels. The backward reactions ranged from a unique lower limit to 0 as an upper bound. And the reversible reactions ranged from -M to M where M is the unique value obtained for each reaction. Hence, GSM models for healthy AM, M1, and M2 phases were obtained by implication of both approaches. We compared iMAT and E-flux algorithms and the details are discussed in the Results and Discussion section. To ensure the biological relevance of these GSM models, we used techniques such as Flux Balance Analysis (FBA)(36) and Flux Variability Analysis (FVA)(37) to analyze and improve model connectivity.

### Flux Balance and Flux Variability Analysis

Flux Balance Analysis (FBA) is used in this study to analyze the flow of metabolites in different conditions. FBA is a widely used approach to study biochemical networks, namely, genome-scale metabolic models that contain the known metabolic reactions in a biological system and the genes that encode each enzyme(36). The GSM model is represented by a stoichiometric matrix which contains metabolites as columns and the rows are represented by reactions. The upper and lower bounds act as a constraint on each of the reactions based on nutrient availability and other microenvironment conditions. FBA generates a flux value for each reaction. Flux Variability Analysis (FVA) is an extension of FBA which calculates the maximum and minimum possible flux for all the reactions in the model at a specific condition(37).

### Model Curation

The three metabolic models were curated by using the classic design-build-test-refine cycle to be able to ensure proper network connectivity and accurate reflection of the metabolic capabilities of the alveolar macrophage cell. Despite the generated models being capable of producing biomass, key metabolite productions such as NO, succinate, and itaconate were found to be different than what was expected in this cell. This could be due to the presence of thermodynamically infeasible cycles (TICs). TICs are cycles created by reactions that carry fluxes even in the absence of nutrients essential for cellular growth and functionality. The TICs can cause the metabolic model to produce metabolites higher/lower than expected, by activating reactions that would be off in a biological scenario. However, if essential reactions are eliminated or the directionality of these reactions are changed without proper review, the behavior of the metabolic model might shift away from the known biological phenomena of the cell. Hence, it is extremely important to refine metabolic models by using efficient and effective methods.

We used OptExpand (inhouse tool, currently unpublished), that has been developed as an expansion upon OptFill, a tool previously developed by our group with different functionalities(100). The initial function of OptExpand was to refine GSMs by removing TICs; however, the process of removing TICs from GSMs was found to be much more difficult than the process of incorporating reactions without creating TICs, and thus the method was upgraded to be able to expand a minimal model i.e., minimum number of biochemical reactions required to satisfy the objective, in our models the number was found to be 143) by adding reactions from a database (the database consisted of all but these 143 reactions from Human1). OptExpand generated three possible solutions to avoid formation of any TICs and ensure optimal connectivity. These solutions consisted of either blocking a reaction completely or changing the direction of the reaction. Before incorporating any changes, an exhaustive literature search was conducted to ensure that none of the biologically relevant pathways were omitted fully or partially affected due to these changes. Special attention was given to novel AM pathways such as production of NO from arginine in healthy AM, production of succinate, itaconate and citrate in TCA cycle during M1 phase and citrulline and urea production in NO cycle during M2 phase. FBA and FVA techniques are used to check the fluxes of the metabolic models ensuring proper network connectivity. All the fluxes from FBA and FVA in the absence of nutrients were found to be zero as expected in healthy and activated AM GSM models and in other conditions the fluxes were found to be in accordance with the biological nature of AM.

### Thermodynamic Analysis

Standard Gibbs Free Energy was calculated for reactions using the equilibrator tool(50). Equilibrator is a tool that uses the composition contribution method to calculate the Gibbs free energy of formation at standard conditions. After acquiring the list of standard Gibbs Free Energy, MAX/MIN driving force (MDF) for the pathways of interest was calculated by using the concentration of metabolites ranging from 1 nM to 10 Mm(50). The range was selected due to a lack of specific experimental metabolomics data and literature evidences suggesting that in the context of metabolic reactions occurring in living cells, the metabolite concentration usually ranges from 1 nM to 10 Mm(50). The MDF analysis was performed with the specified metabolic concentration range to obtain change in Gibbs free energy (ΔG) at different temperatures. The value of Gibbs free energy was further used to calculate the enzyme turnover rate and enzyme saturation (K).

The maximum possible flux obtained from FVA was used to calculate the enzyme turnover rates for the reactions of interest. We compared the Kcat values obtained from DLKcat(101) and SABIO-RK(55) and found modest agreement at best. To this end, we put together elements of a new method (to be deployed in larger scale soon) which is capable of reliable Kcat prediction by explicit molecular modeling of respective enzyme structures, and phylogenetic closeness quantification with other enzymes (with the same EC number) but with reported experimental Kcat measurements from SABIO-RK. Each of these enzyme structures for this study was predicted using geometric deep learning variant structure predictor. These structures were pairwise-similarity matched (using TM-Align) against all other enzymes of the similary family that have reported Kcat in SABIO-RK. These similarity scores were used as a weighting term to ascertain the degree of kinship on the Kcat value of the target enzyme at hand. The inclusion of these bio-aware parameters has allowed us to have high confidence in the Kcat value obtained, which would be missing if we just used a sequence-based Kcat predictor instead. The steps and details regarding the whole protocol for Kcat calculation can be found in supplementary files. The relationship between the Kcat, and K was determined for four different enzymes with two Vmax values obtained from FVA (M1 and M2 phase). The equation(102) used is mentioned below:

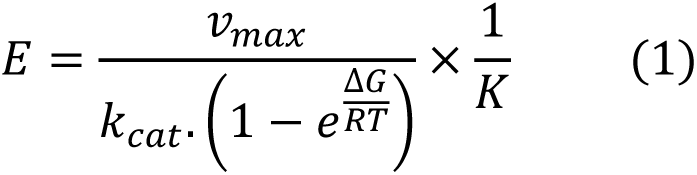

### Structure Informed Kcat Prediction (SI Kcat)

The sequential protocol followed for kcat prediction (illustrated in Figure 8) was demonstrated in one of our most recent works (103) which starts with the data retrieval from SABIO-RK database (43) using specific identifiers such as E.C Number, KEGG Reaction ID, and KEGG Compound IDs. The 3D structures of enzymes are predicted from protein sequences using the RGN2 algorithm, while simultaneously collecting experimentally resolved structures from RCSB PDB (45). Structural comparisons are then made between predicted and experimental structures to assess their similarities (46,47). Utilizing a weighted approach, Kcat values are predicted by considering both structural similarity (Sw) and Kcat data from SABIO-RK. To gauge the uncertainty in predicted Kcat values, pairwise protein sequence alignment is employed.

**Figure 8:**
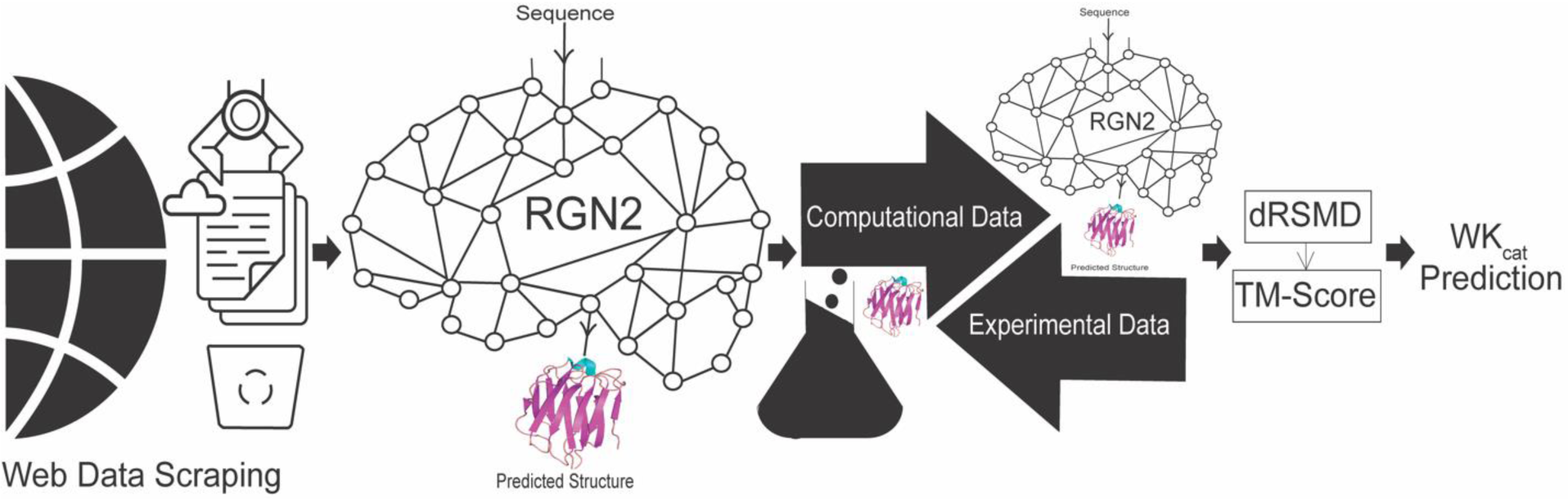
Schematic representation of SI Kcat prediction methodology. The protocol starts with data scraping for Kcat values belonging to the same EC number as the target enzyme, followed by structure prediction for the target sequence using RGN2. The predicted structure is compared with experimental structures for structural similarity weightage (Sw). Si Kcat is calculated following equation 2.

### Measure of Similarity/Dissimilarity between GSM models

The list of reactions that could in any capacity insinuate the metabolic shift from one phenotype to another was deduced by targeting the reactions that displayed distinct flux ranges in both the phenotypes. For example, we started with the list of reactions which had no overlap and definite increase/decrease in forward or backward direction and added some major known metabolic traits. Constraining some reaction fluxes in combination of relaxing certain others could allow the shift of the metabolic fluxes from M2 from M1.

To measure the overall impact and the level of shift upon the inclusion of the constraints added in the GSM models, we explored methods such as Hausdorff distance, Jaccard distance, and Jaccard index. Jaccard index calculates the value based on the intersection and union of a single point data that can be obtained via Eflux2(104). FBA is used to obtain an allowable metabolic flux distribution in a steady-state system in a GSM model, but the obtained fluxes are not unique solutions. We know the GSM models are in general underdetermined, context-specific, and physiologically meaningful flux solutions that can be narrowed down to a unique solution by introducing additional constraints(105). Eflux2 is an extension of FBA that infers a metabolic flux distribution from transcriptomics data and overcomes the shortcoming of E-flux by providing a unique solution. By using this unique solution, the Jaccard similarity index was calculated. Jaccard similarity index is a measure of similarity between two sets of data ranging from 0% to 100%, where the higher percentage indicates higher similarity(106). However, the unique solution of E-flux2 changes with the change in the set value of the objective function. For example, the solution set obtained with the maximum biomass obtained from FBA is different from the solution with the biomass set as the maximum flux from FVA. Since there is no definite growth rate for alveolar macrophage reported to the extent of our knowledge. Hence, the flux ranges from FVA were used for the calculation of Jaccard distance and Hausdorff distance. The additional information including formulation and calculation of Jaccard index, Jaccard distance and Hausdorff distance is available in the supplementary files.

## Data Availability

The GSM models and other supporting files can be found in this GitHub repository: https://github.com/ssbio/Alveolar-Macrophage. The codes for calculation and determination of SI-kcat are available in this GitHub directory: https://github.com/ChowdhuryRatul/kcat_iZMA6517.

## Acknowledgment

We would also like to thank the Holland Computing Center (HCC) of the University of Nebraska, which receives support from the Nebraska Research Initiative (United States of America).

## Author contributions

S.M. and R.S. worked on concept development for this work and developed the methodologies. The analysis followed by the writing was done by S.M. under the guidance of R.S. Kcat predictions were completed by K.A.S., and B.A. under R.C’s supervision.

## Supporting Information

**S1 file:** List of reactions, metabolites and gene associations in the generated GSM models of healthy alveolar macrophage, M1 phase, and M2 phase in excel format.

**S2 file:** SBML file for healthy alveolar macrophage GSM model.

**S3 file:** SBML file for M1 phase GSM model.

**S4 file:** SBML file for M2 phase GSM model.

**S5 file:** SBML file for modified M2GSM model.

**S6 file:** Excel file including the details on thermodynamic parameters calculations such as relationship between Kcar and E.

**S7 file:** Excel file containing details regarding the constraints on the reactions that allow the switch of M2 to M1 phenotype.

